# Fast Hyperspectral and Super-Resolved Mapping of Lipid Membrane Polarity with Single-Molecule Sensitivity

**DOI:** 10.1101/2025.09.19.677386

**Authors:** Elric Dion Pott, Meek Yang, James Ethan Batey, Joie Embree, Bin Dong

**Affiliations:** Department of Chemistry and Biochemistry, University of Arkansas, Fayetteville, Arkansas 72701, United States

## Abstract

Cell membranes display nanoscale heterogeneity in lipid composition and organization that regulates vital biological processes yet remains challenging to resolve with conventional imaging. We introduce spectral phasor single molecule localization microscopy (SP-SMLM), a hyperspectral and super-resolution method that combines wavefront-like optical filtering with single molecule imaging for simultaneous spatial and spectral analysis. A lab-built three-channel imager with sine/cosine filters encodes emission spectra of single molecules into the phasor space, enabling high-throughput, high-SNR mapping of membrane polarity at sub-50 nm spatial and 15-sec temporal resolutions. Through simulation, we validate that the phasor angle correlates with the spectral mean for single dye molecules. When applied to Nile Red-stained COS-7 cells, SP-SMLM revealed organelle-specific polarity differences and dynamic remodeling of lipid composition within live cells. The method’s hyperspectral capability, rapid acquisition, and compatibility with 2D/3D imaging platforms position SP-SMLM as a powerful tool for studying membrane heterogeneity and dynamics in live cells.

## Introduction

Cell membranes are composed of hundreds of distinct lipid species asymmetrically distributed across two leaflets, alongside a multitude of proteins. Historically, membrane biology emphasizes proteins as the principal drivers of membrane functionality and suggests that lipids only play a passive structural role. The lipid raft hypothesis, introduced in 1997, challenged this notion by proposing that specific sphingolipid–cholesterol–protein assemblies contribute actively to membrane trafficking and signaling.^1, 2^ These rafts are characterized by tight lipid packing and chemical specificity, forming nanoscale, sterol-enriched, ordered domains.^3^ Their dynamic and metastable nature allows for coalescence into larger functional assemblies under specific molecular interactions.^4, 5^ Understanding lipid membrane heterogeneity is therefore essential, since it reveals how such domains organize cellular processes and respond to physiological stimuli. Yet, conventional imaging techniques often lack the spatial resolution needed to visualize these nanodomains,^6^ creating barriers to uncovering their roles in membrane organization, composition, and morphology.^6–8^

To overcome the spatial resolution barrier preventing the visualization of lipid membrane heterogeneity, various techniques like structured illumination microscopy (SIM),^9^ DNA-point accumulation for imaging in nanoscale topography (DNA-PAINT),^10^ and spectrally resolved stochastic reconstruction microscopy (SR-STORM)^11, 12^ have been used each with different strengths and weaknesses. SIM enables rapid imaging with a resolution of around 100 nm; however, the resolution is still not high enough to resolve the lipid nanodomains. DNA-PAINT has the spatial resolution to resolve structures at the nanometer scale but suffers from low temporal resolution.^13^ SR-STORM offers simultaneous spectral analysis and spatial localization of single molecules, enabling compositional mapping of lipid domains with high precision; yet it requires a complex imaging system since the spectral and spatial channels need to be precisely aligned. Furthermore, SR-STORM is limited by photon budget and low emitter density due to spectral overlap.^10, 14^

Here, we developed a method for fast hyperspectral super-resolution imaging with single molecule sensitivity. The developed imaging system integrates a non-dispersive spectroscopy measurement with single molecule photoswitching for simultaneously quantifying the spectral color and spatial location of single molecules. The non-dispersive spectroscopy analysis was achieved using lab-built three-channel imager within which the emission signal from single molecules was modulated by two wavefront-like transmission filters, namely, cosine/sine filters. The cosine/sine filters serve as the physical Fourier transformers and convert the emission spectra from single molecules into phasor space with which the spectral information was quantified. We therefore name the method as spectral phasor single molecule localization microscopy (SP-SMLM).

SP-SMLM has unique advantages including single molecule imaging with high signal to noise ratio (SNR), low probability in spectral interference of nearby single molecules, and compatible with dense sample labeling conditions. These unique features of SP-SMLM thus enable high throughput single molecule spectroscopy analysis and hyperspectral super-resolution microscopy imaging with high temporal resolution. We applied the developed method to study the nanoscale heterogeneity in subcellular lipid membranes in both fixed and live cells using a solvatochromic dye molecule, Nile red. The significant difference in lipid membrane polarity among subcellular organelles were clearly observed. The role of cholesterol in controlling lipid membrane polarity was studied and revealed. We also uncovered the dynamics of lipid membrane polarity in live cells, which reflects the highly exchangeable feature of lipid compositions. Being able to map lipid membrane polarity could allow us to better understand how cholesterol is trafficked within neurons and how cholesterol is shuttled between the endolysosomal system, endoplasmic reticulum, and mitochondria.^15^ This could give us new tools for studying neurodegenerative diseases, since it is known that alterations in cholesterol homeostasis are associated with disruption of brain function.^16–19^

## Results and Discussions

### SP-SMLM enables simultaneous spectroscopy analysis and spatial localization of single emitters at high density

The schematic view of the SP-SMLM imaging system is shown in Figure 1a (Supporting Information S1, Figure S1). To quantify spectral information of the collected fluorescence signal from single emitters, it was directed into a lab-built three-channel imager. The emission signal was first split by a 30:70 nonpolarizing beam splitter. The transmitted portion (i.e., 70%) was further split by another 50:50 nonpolarizing beam splitter. Two additional optical wide-transmission bandpass filters (i.e., cosine and sine filter, Figure 1b) were inserted into the two optical paths from the 50:50 beam splitter. All three optical paths were directed to a highly sensitive electron-multiplying charge-coupled device (EMCCD) camera using a pair of relay lens. As a result, three images (namely, cos, sin, and ref) of the same single emitters (Figure 1c-e) are simultaneously captured. The relative photon intensities in the three channels depend on the spectral property (i.e., color and width) of single emitters (Figure S2-3, Supporting Information S2). Therefore, one can obtain spectral information of single emitters without the need of any dispersive optics.

**Figure 1.**
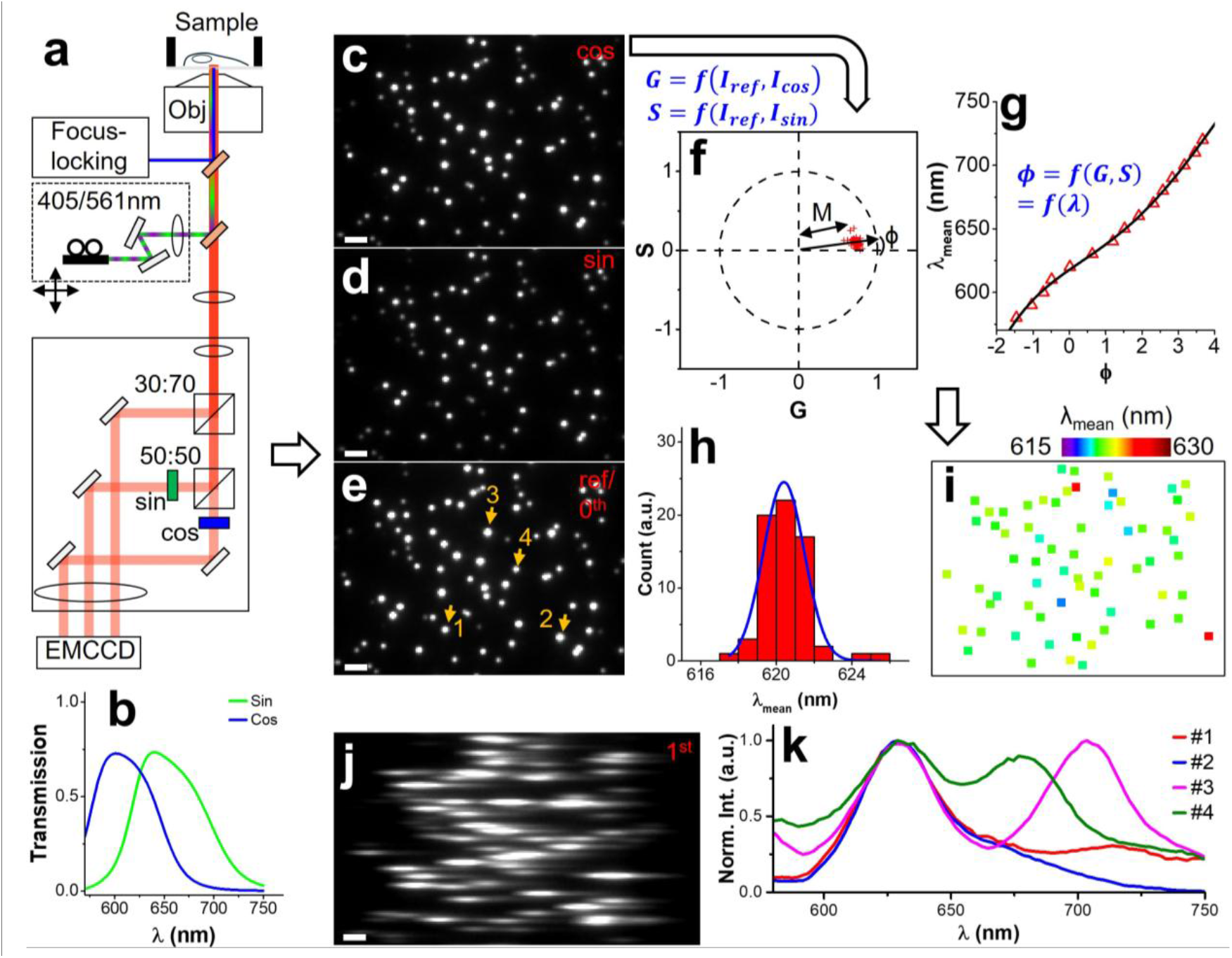
Working principle of high throughput SP-SMLM. (a) Schematic of the optical setup. Collected emission signals from single emitters are separated into three channels with two of them modified by sine and cosine optical filters. (b) Transmission profiles of sine and cosine filters. (c-e) cos-, sin-modified and nonmodified (i.e., ref) or 0^th^ order diffraction image of single fluorescent beads. (f) Phasor plot of fluorescent beads as single emitters. (g) Calibrated wavelength-phase angle relationship. (h) Histogram distribution of calculated spectral mean of single fluorescent beads. (i) Hyperspectral SMLM images of single fluorescent beads. (j) 1st order diffraction image of the same fluorescent beads in (e). (k) Examples of fluorescence emission spectra from single fluorescent beads. Significant spectral interference was observed under high emitter density (#1, 3-4). Scale bar: 2 µm.

In SP-SMLM, the same single emitters in the three channels (Figure 1c-e) were first localized and matched. Their photon intensities in cosine, sine, and reference channels were used to calculate single emitters’ locations in the phasor space (i.e., G and S, Figure 1f, Supporting Information S3). In the phasor space, one can obtain two important parameters, i.e., phase angle (ϕ) and amplitude (M), which are correlated to the spectral mean (λ_mean_) and width of single emitters respectively (Figure S2-3, Supporting Information S2). Since there is no dispersion in single emitters’ images, this approach is free of spectral interference at densities up to diffraction limit. To build the correlation between phase angle and spectral color, we calibrated our imaging system using the light source of a microscope lamp. Brightfield images of a clean sample slide were captured at different wavelengths (Figure S4) by inserting narrow bandpass filters into the optical path (Supporting Information S4). Based on these images, a phasor plot was then constructed to determine the phase angle at each wavelength. The resulting wavelength-phase angle relationship (Figure 1g) was well-fitted by a fourth-order polynomial function. This calibration curve was then used to determine the spectral color of single emitters (Figure 1h). The intensity-weighted spatial locations from three channels were calculated and used for super-resolution image reconstruction. A hyperspectral SMLM image was reconstructed where the color represents the spectral mean of single emitters (Figure 1i). Although both the spectral mean and width of emission signal can be determined from the phasor plot. In this work, we focused on the quantification of spectral color of single emitters.

In comparison, the spectral interference from nearby emitters was readily present using the dispersive method (Figure 1j-k, Figure S5). This poses a challenge for accurate spectroscopy analysis at high density of single emitters. Furthermore, using a non-dispersive approach also results in images with high SNR which reduce uncertainties in both spatial locations and spectral measurements. Moreover, photons from all channels were combined to determine the spatial locations of single emitters thus improving the spatial resolution in SP-SMLM. Therefore, SP-SMLM also has better photon usage. Overall, these advantages of SP-SMLM set its potential for image acquisition at a fast rate, high SNR and emitter density. Therefore, hyperspectral super-resolution imaging at high spatial and temporal resolutions can be achieved.

### SP-STORM reveals heterogeneity in lipid membrane polarity at subcellular organelles

In mammalian cells, lipid membranes compose glycerophospholipid, sphingolipid, and cholesterol with decreased polarity respectively.^20^ Nile red (NR), a membrane permeable and solvatochromic dye shows a blue shift in its emission spectra as the environmental polarity decreases (Figure S6).^20^ Therefore, Nile red has often been used for probing the heterogeneity of lipid membranes’ composition across subcellular organelles.^9, 10, 12^ Additionally, NR is photoswtichable and had been previously used for super-resolution microscopy imaging in direct stochastic optical reconstruction microscopy (dSTORM).^21^ In this work, we applied the concept of SP-SMLM for dSTORM imaging (i.e., SP-STORM) of NR stained cells.

We fixed and labeled the lipid membranes of COS-7 cells with NR (Supporting Information S6). Upon illumination with a high-power density laser, the majority of NR molecules were photoswichted into a nonfluorescent dark state, leaving a random subset and spatially resolved single NR molecules fluorescing (on state) in each imaging frame (Figure 2a). Single NR molecules in all three channels were localized and the corresponding molecules were matched using the method developed in our previous work.^22^ The intensity-weighted locations for single NR molecules were calculated from the individual localized positions in three channels and then used for super-resolution image reconstruction. The same single NR molecules’ photon intensities in three channels were used to determine their phase angle in the phasor plot and their spectral color using the established calibration curved described above. Finally, hyperspectral super-resolved mapping of lipid membrane of subcellular organelles was reconstructed where the color denoting the spectral mean of single NR molecules (Figure 2b). Since the spectral color of NR depends on the polarity of lipid environment. The color in the dSTORM image also represents the polarity of lipid membranes.

**Figure 2.**
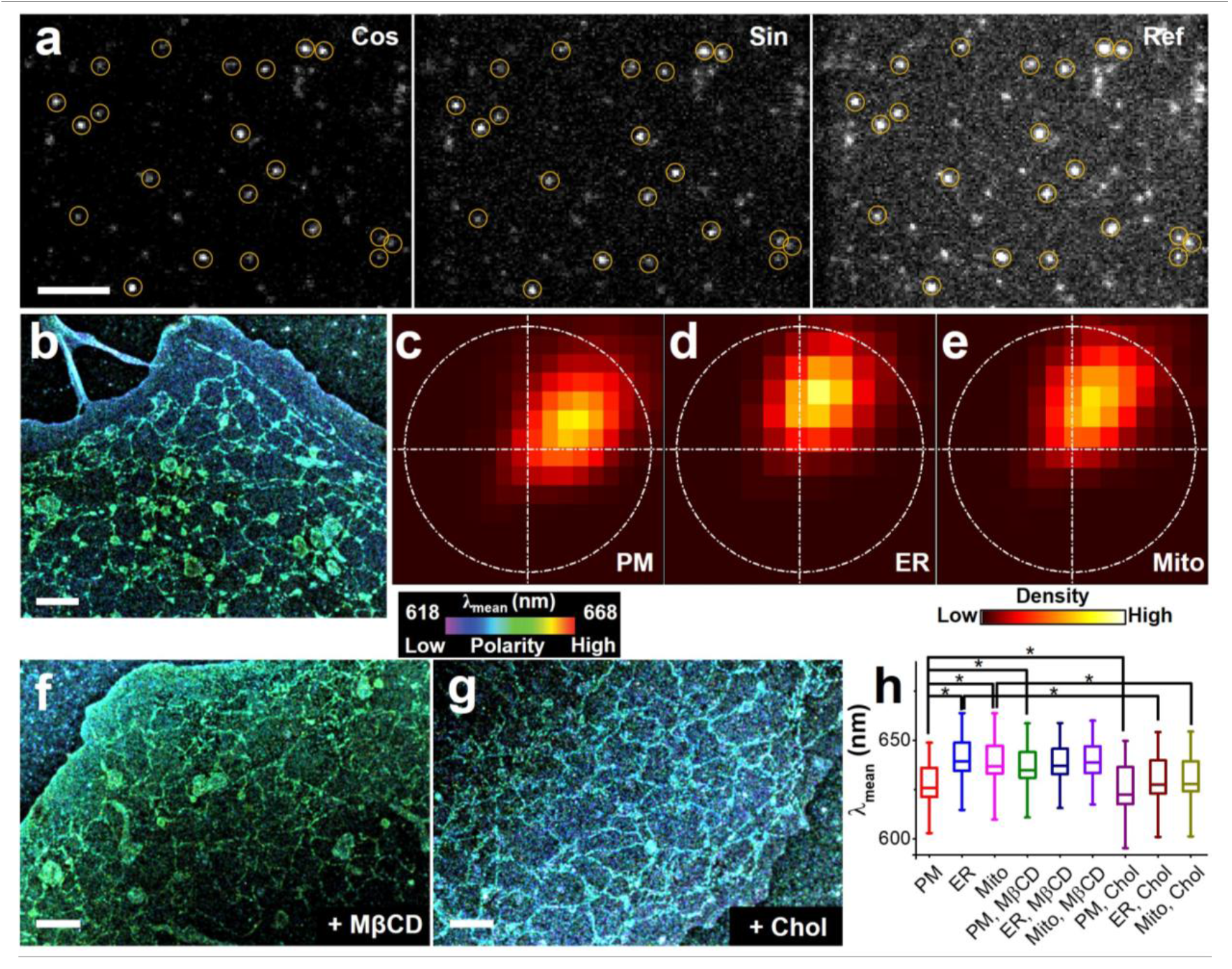
Super-resolved mapping of heterogeneity in lipid membranes by SP-STORM. (a) A snapshot of single molecule NR molecules in cos-, sin-, and ref-channels in SP-STORM imaging. Same single NR molecules are highlighted in yellow circle. (b) Hyperspectral dSTORM image of NR labeled fixed COS-7 cells. (b-d) Phasor plot of single NR molecules at PM, ER membrane, and Mito membrane in (b). Hyperspectral dSTORM image of NR labeled fixed COS-7 cells after depletion of cholesterol with 5 mM MβCD (f), and with incubation of 5 mM water soluble cholesterol (g). (h) Distributions of spectral mean of single NR molecules at PM, ER membrane, and Mito membrane in fixed COS-7 cells under different treatment conditions. Number of single NR molecules, n>10^3^. *: two-tail t-test with p < 0.001. Scale bar: 5 µm (a), 2 µm (b, f, g).

The morphology and structure of subcellular organelles can be clearly distinguished. Furthermore, distinct colors were observed for plasma membrane (PM), endoplasmic reticulum (ER), and mitochondria (Mito) indicating heterogeneity in lipid polarity and compositions across subcellular organelles. We selected three regions of interest (ROIs) that correspond to PM, ER, and Mito. The phasor plots of single NR molecules in these ROIs are shown in Figure 2c-e. The phase angle of single NR molecules in PM is clearly smaller than those in ER and Mito indicating the shorter emission wavelength of NR in PM. These results suggest that the lipid membranes of PM are less polar than those of ER and Mito. We further treated the cells with 5 mM Methyl-β-cyclodextrin (MβCD) and water-soluble cholesterol for the depletion and enrichment of cholesterol in cells respectively (Figure 2f, g). Significant changes in polarity were observed for three types of lipid membranes under both treatment conditions (Fig. 2h). Results with depletion and addition of cholesterol suggest that cholesterol is the driving force behind the membrane polarity differences.

### SP-STORM reveals highly dynamic structural change and lipid compositions exchange of endoplasmic reticulum

In live cells, lipid membranes’ composition, organization, and morphology change constantly, which is essential for regulating the biophysical membrane properties, membrane protein functionalities, and lipid–protein interactions.^23^ These biological processes are highly dynamic ranging from minutes to microseconds.^24, 25^ The high throughput feature of SP-STORM has the advantage of investigating these highly dynamic biological processes. We applied SP-STORM to study the heterogeneity and dynamic regulation of endoplasmic reticulum (ER) in live cells using NR dye. Within a few minutes, >10^6^ single NR molecules were collected (Supporting Movie 1) and used to reconstruct the hyperspectral dSTORM image where the color denotes the spectral mean of single NR molecules (Figure 3a). The spatial resolution of SP-STORM imaging using NR dye in live cells was estimated to be ∼47 nm by small cluster analysis (Figure S7). We obtained hyperspectral dSTORM images in 15 seconds for the ER with localization densities that correspond to a Nyquist spatial resolution of 55 ± 4 nm (s.d., n = 20, Supporting Information S3) for 10 snapshots (Figure 3b, Supporting Movie 2). Dynamic remodeling of ER (e.g., extending tubules, retracting tubules, swing tubules, and extending sheets) were clearly observed. More excitingly, the heterogeneity in polarity within ER tubules (i.e., heterogeneous lipid compositions) and its dynamic changes over time periods were also readily observed by SP-STORM imaging (Figure 3c). The heterogeneous lipid polarity along a single ER tubule over the entire time periods were plotted in Figure 3d. The level of heterogeneity in lipid polarity was revealed to be more than 3x larger under 15-sec temporal resolution than that of 150 seconds. Figure 3e shows the dynamic change of membrane polarity at two close locations on the same ER tubule indicating the exchange of lipid contents over time. Overall, these results uncovered the highly dynamic feature of lipid membrane highlighting the advantages of high temporal resolution in SP-SMLM imaging.

**Figure 3.**
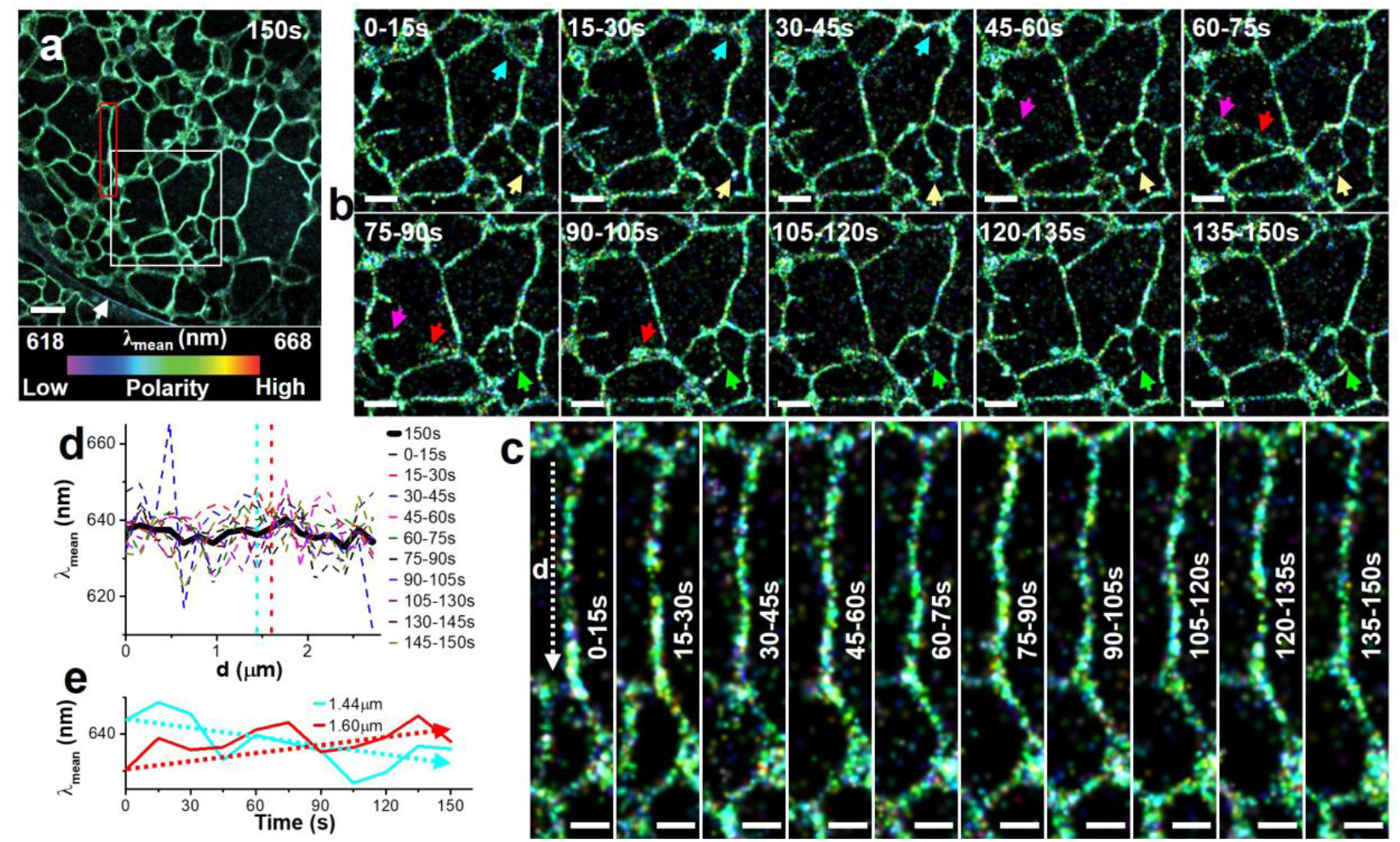
Live cell super-resolved mapping of heterogeneity in lipid membranes by SP-STORM. (a) Hyperspectral dSTORM image of a NR labeled live COS-7 cell over 150 s data acquisition. (b, c) A time-series of 15-sec hyperspectral dSTORM image snapshots of ER dynamics corresponding the white and red boxes in (a). Arrows highlights the dynamic remodeling of ER meshwork in (b). (d) Heterogeneity in lipid membrane polarity along an ER tubule (white arrow in (c)) and its dynamic changes over time. (e) Dynamic change of lipid membrane polarity over time at two nearby locations in an ER tubule. Scale bar: 2 µm (a), 1 µm (b) and 500 nm (c).

## Conclusions

In this work, we developed a high throughput single molecule spectroscopy and hyperspectral single molecule localization microscopy based on non-dispersive spectroscopy analysis approach, i.e., SP-SMLM. Using SP-SMLM, we revealed the heterogeneity of lipid membranes across subcellular organelles and the role of cholesterol as potential driving forces in membrane polarity differences. We also demonstrated SP-SMLM’s applicability in studying live cell dynamics by super-resolved mapping of the dynamic regulation of lipid membranes in ER. The heterogeneity and dynamic change of membrane polarity were observed within the intricate meshwork of ER tubules and sheets, which reflects the highly dynamic nature in lipid composition exchange within the ER meshwork. Further improvement of the throughput of SP-SMLM could allow even faster dynamic processes such as mitochondrial fission and fusion intermediates to be studied. Single-color dSTORM imaging of membrane dynamics with temporal resolution of 1-2 s^24^ had been achieved by boosting high density of single molecules per imaging frame and recording imaging data at higher speed (∼kHz). Similar strategies can be applied for SP-SMLM imaging in future work.

## Acknowledgments

This work was supported by the Department of Chemistry at University of Arkansas Fayetteville, and Starter Grant from the Society for Analytical Chemists of Pittsburgh (SACP).

