## Supporting Information for "Fast Hyperspectral and Super-Resolved Mapping of Lipid Membrane Polarity with Single-Molecule Sensitivity"

**S1. The imaging system.**

The SP-SMLM was carried out on an Olympus IX-81 inverted microscope (Supplementary Fig. 1) equipped with an oil immersion objective (UPLAPO100X, NA 1.50, Olympus). 405 nm (Coherent) and 561 nm (Coherent) lasers were coupled into an optical fiber, recollimated by a fiber coupler, and focused on the back focal plane of the oil immersion objective by a lens after passing a pair of relay lens. Lasers were directed into the sample by a dichroic mirror (ZT561rdc, Chroma). All Lasers were cleaned by a multi-bandpass filter. A translation stage was used for shifting the laser beams laterally before entering the objective so that the laser beams project to the sample/coverslip interface at an angle slightly smaller than the critical angle. The emission signals from dye molecules were collected by the same oil immersion objective. A long pass emission filter (ET575lp, Chroma) was used to reject laser scattering background. Then, the collected signal was directed to a lab-built three-channel imager for in-hardware Fourier transformation based spectral phasor analysis. Inside the three-channel imager, the emission signal from dye molecules was first split by a 30:70 (R:T) nonpolarized beam splitter (BS019, Thorlabs). The transmitted light after the first beam splitter was split further by a 50:50 (R:T) nonpolarized beam splitter (BS013, Thorlabs). The reflected and transmitted signals from the second beam splitter were transformed by sine and cosine function-like optical filters respectively. Three images of the same dye molecules, i.e., reference, sin/cos-modified images, were projected on different regions of the same electron-multiplying charge-coupled device (EMCCD) camera (iXon Ultra 897, Andor) using a pair of relay lens. A short pass optical filter (FESH0750, Thorlabs) was used to confined spectral window below 750 nm. An optical slit was placed at the intermediate image plane of the tube lens allowing crop field of view and avoiding spatial overlap between three channels. The sample stage was locked by a self-built autofocusing optical system running by self-written Python scripts.

**S2. Simulation study.**

The locations of dye molecules in the spectral phasor plot depend on both the wavelength and the width of emission spectra, which lays the foundation for spectral demixing in phasor analysis. We use simulation results to demonstrate the working principle of spectral demixing based on phasor analysis and the effects of spectral properties on the performance. We simulated the fluorescent emission spectra at six peak wavelengths (15 nm spectral interval) and four peak widths (full width at half maximum, FWHM= ~2.355σ: 10, 20, 50, and 100 nm) using the normalized Gaussian equation:

$$I(\lambda) = \exp\left(\frac{-1 * (\lambda - \lambda_{peak})^2}{2\sigma^2}\right)$$

where I(λ), $\lambda_{peak}$, and σ are intensity at λ, the peak wavelength, and standard deviation of the peak, respectively. The results are shown in Figure S2-S3.

*S2.1. Using ideal sine/cosine transmission filters.* Spectral phasor analysis using ideal sine/cosine filters were first conducted. The ideal sine/cosine transmission filters are simulated using the following equations:

$$T_{cos}(\lambda) = \cos\left[2\pi\left(\frac{\lambda}{\lambda_{max} - \lambda_{min}} + n\right)\right]$$

$$T_{sin}(\lambda) = \sin\left[2\pi(\frac{\lambda}{\lambda_{max} - \lambda_{min}} + n)\right]$$

$$n = 1 - \frac{\lambda_{min}}{\lambda_{max} - \lambda_{min}}$$

where $T_{sin}(\lambda)$ and $T_{cos}(\lambda)$ are the transmission efficiency for sine and cosine filter at $\lambda$; $\lambda_{min}$ and $\lambda_{max}$ are the low and high end of detection window of wavelength; n is the correction factor that enables one period of sine/cosine wavefunction within the [$\lambda_{min}$, $\lambda_{max}$]. The simulated transmission profiles of ideal sine/cosine filters are shown in Figure S2. Applying the sine/cosine transmission filters on the simulated emission spectra transforms them into two modified emission spectra (Figure S2b-c). The phasor plot (G, S) can then be constructed using the following equation:

$$G = 2\frac{\int_{\lambda_{min}}^{\lambda_{max}} I(\lambda)\otimes T_{cos}(\lambda)d\lambda}{\int_{\lambda_{min}}^{\lambda_{max}} I(\lambda)d\lambda} - 1$$

$$S = 2\frac{\int_{\lambda_{min}}^{\lambda_{max}} I(\lambda)\otimes T_{sin}(\lambda)d\lambda}{\int_{\lambda_{min}}^{\lambda_{max}} I(\lambda)d\lambda} - 1$$

The results in spectra phasor plot for different peak wavelengths and widths are shown in Figure S2d. The results show two obvious trends: larger phase angles for emission at longer wavelengths and smaller phase amplitudes for wider emission spectra. In this work, the wavelength-phase angle relationship is of the most interest. Two types of wavelength-phase angle relationship were plotted using spectral peak and mean respectively (Figure S2e-f). Spectral mean was calculated as the intensity-weighted averaging of wavelength. Significant variation of peak wavelength-phase angle relationships was observed for wide emission spectra (FWHM > 20 nm) due to the confined boundary of spectral detection window. On the contrary, the relationship between spectral mean and phase angle maintains very well (FWHM < 50 nm). In fact, phasor analysis was developed as a non-fitting method for analyzing spectral data and to obtain the spectral mean.

*S2.2. Using actual sin/cos filters.* We further studied the effects of peak wavelength and width in the spectra phasor results when using the actual sine/cosine transmission filters in our experiments. Figure S3 shows the transmission profiles of the sine/cosine transmission filters measured by a UV-Vis spectrometer. Similarly, transformation of emission spectra with actual sine/cosine transmission filters can be done as that used with ideal sine/cosine transmission filters (Figure S 3b-c). To construct the phasor plot (G, S), the following modified equations were used to accommodate the min/max transmission of the actual sine/cosine transmission filters:

$$G = 2\frac{\frac{\int_{\lambda_{min}}^{\lambda_{max}} I(\lambda)\otimes T_{cos}(\lambda)d\lambda}{\int_{\lambda_{min}}^{\lambda_{max}} I(\lambda)d\lambda} - T_{cos,min}}{T_{cos,max} - T_{cos,min}} - 1$$

$$S = 2\frac{\frac{\int_{\lambda_{min}}^{\lambda_{max}} I(\lambda)\otimes T_{sin}(\lambda)d\lambda}{\int_{\lambda_{min}}^{\lambda_{max}} I(\lambda)d\lambda} - T_{sin,min}}{T_{sin,max} - T_{sin,min}} - 1$$

The results are shown in Figure S3d. A nonlinear relationship between wavelength (either spectral peak or spectral mean) and phase angle were obtained (Figure S3e-f), which matches with the calibration results using mechanical slits and bandpass filters (Figure S4c). The same results were also observed for the variation of wavelength-phase angle relationships. The spectral mean-phase angle relationship maintains the trend for emission spectra with FWHM < 50 nm which is within the range of FWHMs for Nile red dye in different polarity environment.

**S3. Data analysis.**

The collected single molecule imaging data were analyzed by either ThunderStorm[1] ImageJ plugin or Insight3 software. The single molecules in all three channels were first identified and localized independently. To identify the same single molecule in all channels, a previously developed correlation analysis procedure was adapted.[2] We first imaged fluorescent beads on coverslip and use their localized positions to build a transformation matrix. The transformation matrix can then be used to project the spatial coordinates of localized single molecule positions from sine- and cosine-modified channel to the refence channel. The projected molecular positions are then compared to those obtained in the refence channel. Molecular positions that are within the same imaging frame and within the localization precision are considered as from the same molecule. Their photon intensities were used for constructing phasor plot (G, S) and calculating the phase angle (ϕ) using the following equations:

$$G = 2\frac{(\frac{I_{cos}}{fI_0} - T_{\cos,min})}{(T_{\cos,max} - T_{cos,min})} - 1$$

$$S = 2\frac{(\frac{I_{sin}}{fI_0} - T_{\sin,min})}{(T_{\sin,max} - T_{sin,min})} - 1$$

$$\phi = \tan^{-1}(\frac{S}{G})$$

where $I_0$, $I_{sin}$, $I_{cos}$ are photon intensity of single molecules in reference, sine, and cosine channels, respectively. f (~1.17) is the correction factor for the intrinsic difference in transmission efficiency between reference and sine/cosine channels. $T_{cos,max}$ and $T_{cos,min}$ are the maximum and minimum transmission of the cosine filter in the confined spectral window. $T_{sin,max}$ and $T_{sin,min}$ are the maximum and minimum transmission of the sine filter in the confined spectral window. Applying the calibration curve of spectral mean-phase angle relationships, we obtained the spectral mean ($\lambda_{mean}$). The final localized positions of single molecules were intensity-weighted average from those in all channels. Combining the obtained spectral mean and weighted average localized positions of single molecules, hyperspectral dSTORM image was rendered using Insight3 software. In snapshots of live cell dynamics, the Nyquist resolution limit was estimated based on localization densities in SP-STORM images. Small regions of ER tubules were selected to measure the area (A) and to determine the number of localizations (n) within the area of interest. The Nyquist spatial resolution was calculated as $R_s = \frac{2}{\sqrt{\frac{n}{A}}}$. Post-data analysis of localized single molecules was done using MATLAB scripts and Insight3.

**S4. Calibration for wavelength-phase angle relationship in SP-SMLM.**
We calibrated the wavelength-phase angle relationships of the SP-SMLM imaging system using the brightfield images of transmitted light from the microscope lamp. The lamp light was filtered by a series of narrow bandpass filters (Thorlabs) giving three images (i.e., reference, sine-, and cosine-modified) at specific wavelengths (Figure S4a). Measuring the intensities from three channels allows the determination of locations of spectral color in the phasor space (Figure S4b). The calibration curve of phase angle-spectral color was well fitted by a fourth-order polynomial function (Figure S4c).

**S5. Calibration for wavelength-pixel shift relationship in grating based SR-SMLM.**
To calibrate the relationship between wavelength and pixel shift in a gratings-based SR-SMLM system, we introduced a grating into the detection pathway of our microscope, producing both $0^{th}$ and $1^{st}$ order diffraction images (Figure S5a). We illuminated a glass slide without sample using a microscope lamp and filtered the light through a series of narrow bandpass filters to isolate specific wavelengths. For each wavelength, we determined the intensity centers of the $0^{th}$ and $1^{st}$ order images (Figure S5b), calculated the corresponding pixel shifts, and fitted the pixel shift wavelength relationship to a linear model (Figure S5c).

**S6. Sample preparation.**
*S6.1. Cell culture.* COS-7, African green monkey kidney cells (CRL-1651, ATCC) was purchased from the American Type Culture Collection (ATCC). The COS-7 cells were cultured in T25 cell culture flask (690160, Greiner Bio-One) with the cell culture medium Dulbecco's Modified Eagle's Medium (DMEM) (10-014-CV, Corning) added with 10% fetal bovine serum (FBS) (26140079, Gibco) along with 1% Penicillin-Streptomycin (Pen Strep) (15140122, Gibco) (i.e., cell culture media). µ-dish (81158, ibidi) was purchased from ibidi for subculturing cells. To subculture cells in µ-dish, 120 µL of cell suspension solution was added into the dish, followed by 2 mL of complete cell culture media. The cells were allowed to incubate for 24 hours at 37°C and 5% $CO_2$ inside the cell culture incubator before imaging.

*S6.2. Cell labeling with Nile red dye.* To label fixed cells with Nile red, the COS-7 cells were rinsed once with pre-warmed DPBS buffer (21-030-CV, Corning), fixed with pre-warmed 3% (v/v) paraformaldehyde (0433689M, ThermoFisher) + 0.1% (v/v) glutaraldehyde (340855, Sigma) in 1× DPBS buffer for 20 minutes, reduced with freshly prepared 0.3% (m/v) $NaBH_4$ (BDH4604, VWR) for 10 minutes, followed by three times of washes with DPBS buffer. The cells were stained with 100-500 nM Nile red (41517110000, Thermo Scientific) in DPBS buffer for 30 minutes. The cells were briefly washed three times with DPBS buffer before mounting on the microscope for imaging. To label live cells with Nile red, the COS-7 cells were rinsed once with pre-warmed Leibovitz's L-15 medium (10-045-CV, Corning), incubated with 100-500 nM Nile red in L-15 medium for 30 minutes at 37 °C with 5% $CO_2$ inside the cell culture incubator. The cells were briefly washed three times with L-15 medium before mounting on the microscope for imaging.

*S6.3. Cholesterol in addition to cells.* For Cholesterol addition, the cells were treated with a 5 mM water soluble cholesterol (C4951-30MG, Sigma Aldrich) in L-15 medium for live cells or DPBS buffer for fixed cells for 60 minutes. This step was done before the sample was stained with Nile red.

*S6.4. Cholesterol depletion from cells.* For Cholesterol depletion, the cells were treated with 5 mM Methyl-β-cyclodextrin (C4555-1g, Sigma Aldrich) in L-15 medium for live cells or DPBS buffer for fixed cells for 20 minutes. This step was done before the sample was stained with Nile red.

**S7. SP-STORM imaging.**
The dye labeled samples were first excited at low laser power for locating cells and capturing conventional fluorescence images. Laser power was then increased to high power density (1-2 kW $cm^{-2}$) to photoswitch most of the Nile red molecules into nonfluorescent dark state. A random subset and spatially resolved Nile red molecules were kept in fluorescing (on state) at any given instant. 405 nm laser at low power density was used to adjust the density of single Nile red molecules per image frame when it is necessary. The EMCCD camera acquired images in all three channels (i.e., reference, sine- and cosine-modified) simultaneously and continuously at a frame rate of 200 Hz. The imaging data was typically recorded for 30k frames, which corresponds to an acquisition time of 2.5 minutes. SP-STORM imaging of Nile Red was carried out in a 200 µM ascorbic acid (BDH9242, VWR) in L-15 medium for live cell imaging and in DPBS buffer for fixed cell imaging.

### S8. Supporting Figures

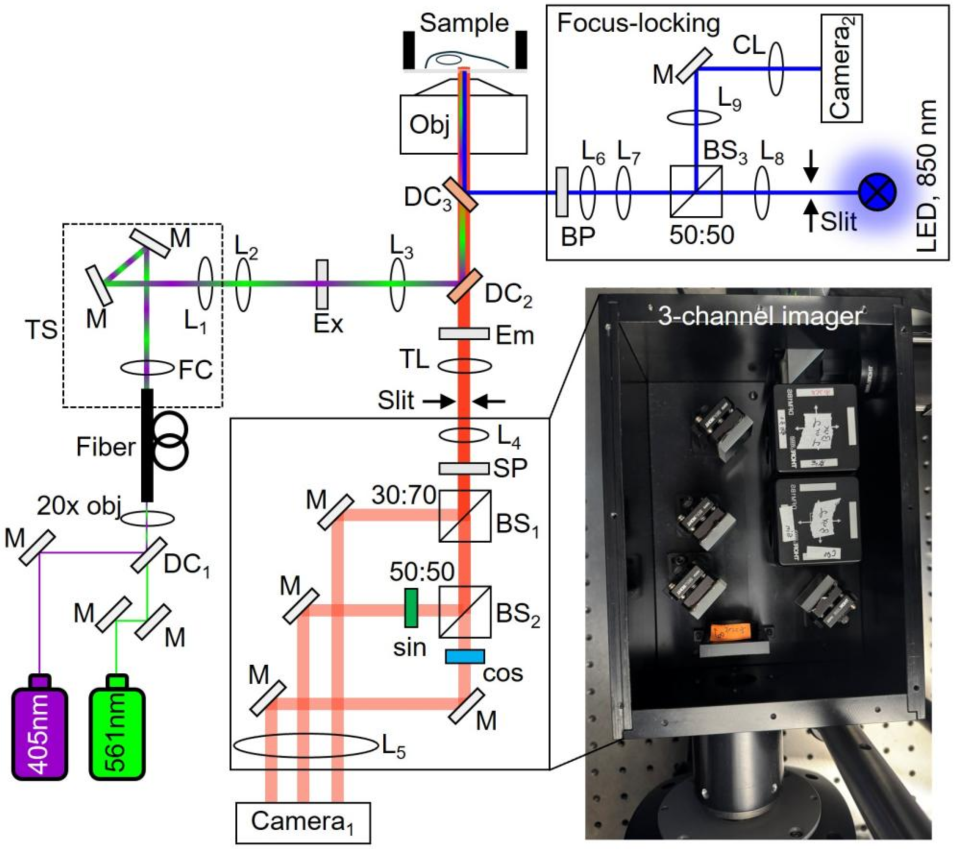

**Figure S1. Optical setup of SP-STORM.** The imaging system is built with an Olympus IX81 motorized inverted microscope.

Light sources: Lasers, 100 mW 405 nm (Coherent Obis), 150 mW 561 nm (Coherent Obis); LED, 1400 mW 850nm (M850LP1, Thorlabs).

Optics: M, BB1-E02 or BBSQ1-E02 or MRA25-E02 (Thorlabs); Lens1-5, AC254-100-A, AC254-100-A, AC254-200-A, AC254-125-A, AC254-125-A (Thorlabs); Lens 6-9, LB1945-AB, LB1676-AB, LB1757-AB, LB1901-AB (Thorlabs); CL, LJ1703RM-B (Thorlabs); DC1, 86-389 (Edmund Optics); DC2, ZT561rdc (Chroma); DC3, FF750-SDi02 (Semrocks); Ex, ZET405/488/561/640xv2 (Chroma); Em, ET575lp (Chroma); SP, FESH0750 (Thorlabs); BS1, BS019 (Thorlabs); BS2, BS013 (Thorlabs); BS3, BS014 (Thorlabs); BP, FF01-857/30 (Semrock); sine/cosine filters, 650FS80/600FS80 (Andover).

Objectives: 20× air objective (Olympus); 100× Oil TIRF objective (UPLAPO100XOHR, N.A. 1.50, Olympus)

Fiber components: FC, TC18APC-543 (Thorlabs); fiber, P3-S405-FC-1 (Thorlabs).

Mechanic and motorized components: TS, PT1-Z9 and KDC101 (Thorlabs); slit, VA100CP (Thorlabs).

Detectors: camera 1, Andor iXonEM+ Ultra 897; camera 2, DCC1545M (Thorlabs).

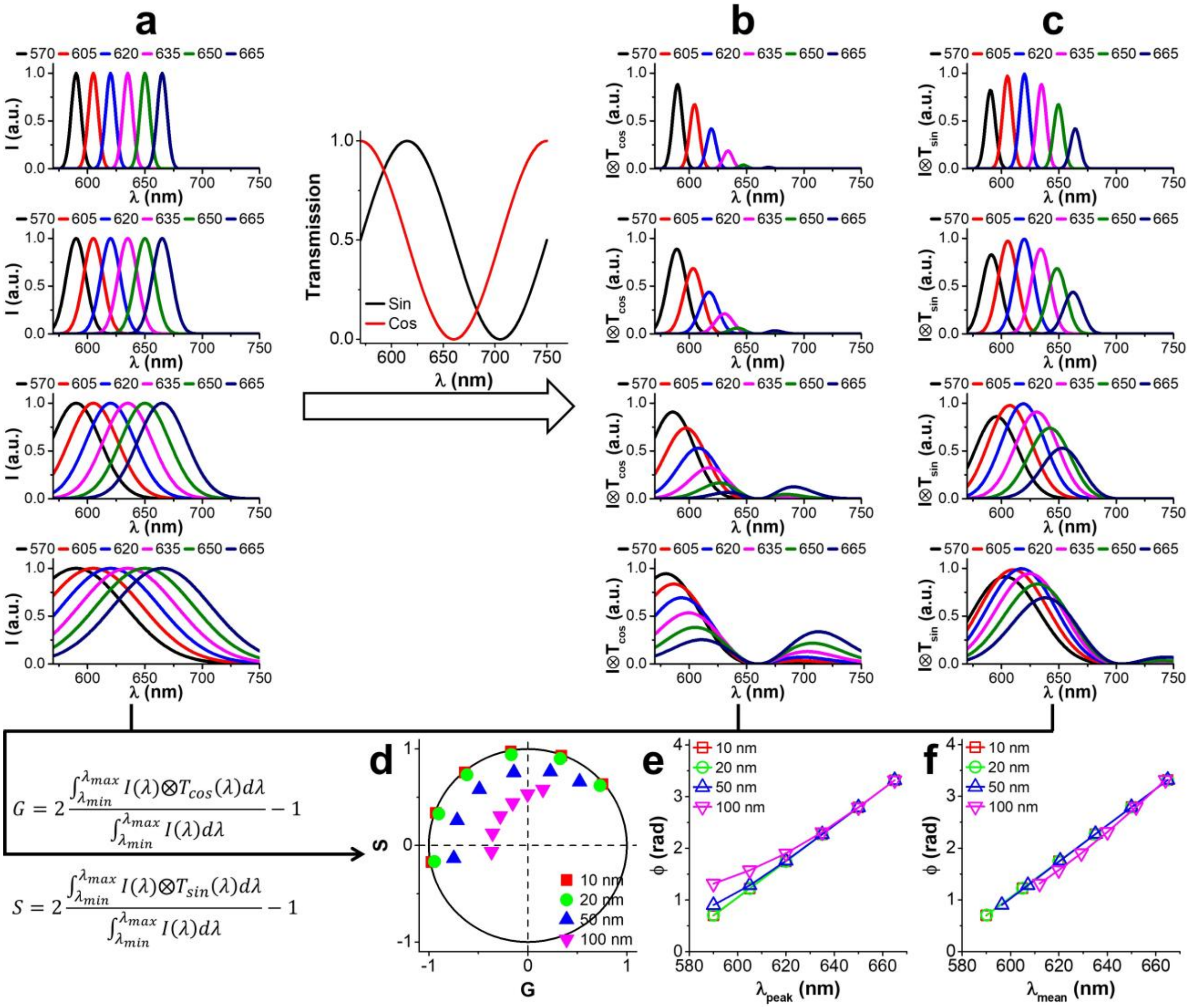

**Figure S2. Spectral phasor analysis with ideal sine and cosine filters.** (a) Simulated emission spectra with Gaussian distribution at different peak wavelengths and widths. (b) Cosine and (c) sine modified emission spectra from (a). (d) Phasor plot. Black solid line denotes results directly calculated from transmission profiles of ideal sine and cosine filters, representing phasor plot at single-wavelength resolution. (e) Phase angle-peak wavelength and (f) phase angle-spectral mean relationships, respectively.

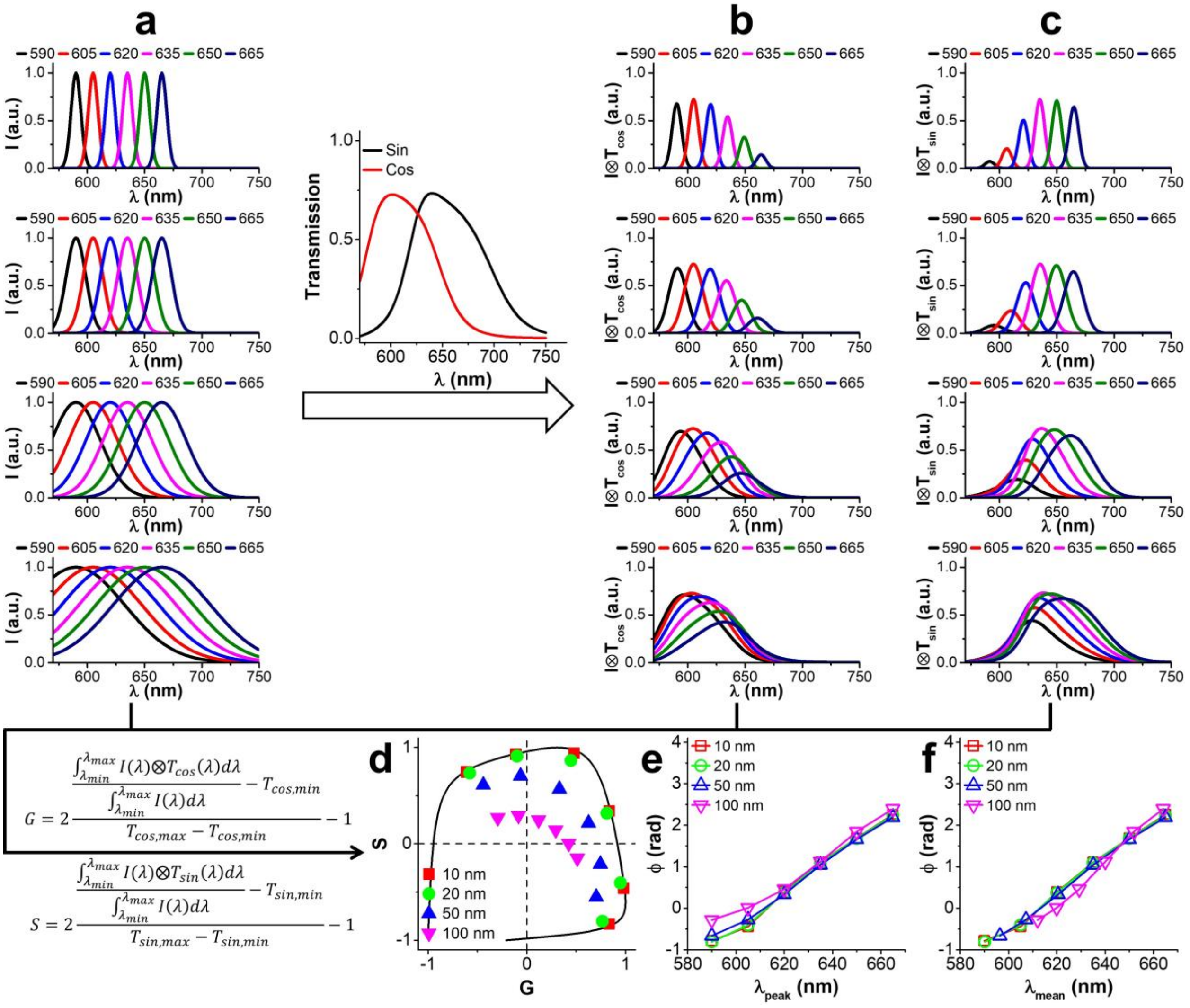

**Figure S3. Spectral phasor analysis with actual sine and cosine filters.** (a) Simulated emission spectra with Gaussian distribution at different peak wavelengths and widths. (b) Cosine and (c) sine modified emission spectra from (a). (d) Phasor plot. Black solid line denotes results directly calculated from transmission profiles of actual sine and cosine filters, representing phasor plot at single-wavelength resolution. (e) Phase angle-peak wavelength and (f) phase angle-spectral mean relationships, respectively.

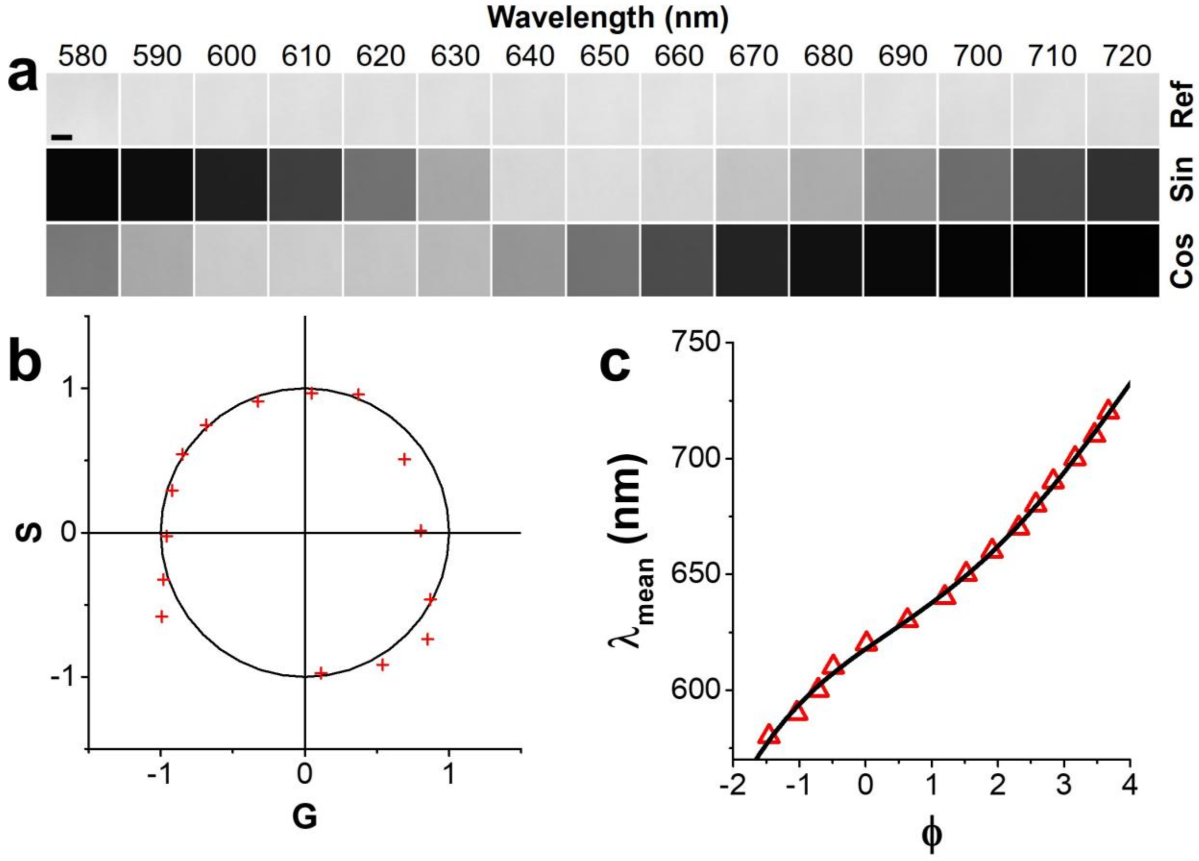

**Figure S4. Calibration of the optical setup of SP-SMLM.** (a) Brightfield image of microscope lamp light at different wavelengths. Narrow bandpass filters of different wavelengths were inserted into the optical path of detection. (b) Phasor plot of results obtained from (a). (c) Calibrated wavelength-phase angle relationship. The data was fitted with a fourth-order polynomial function. Scale bar: 5 μm.

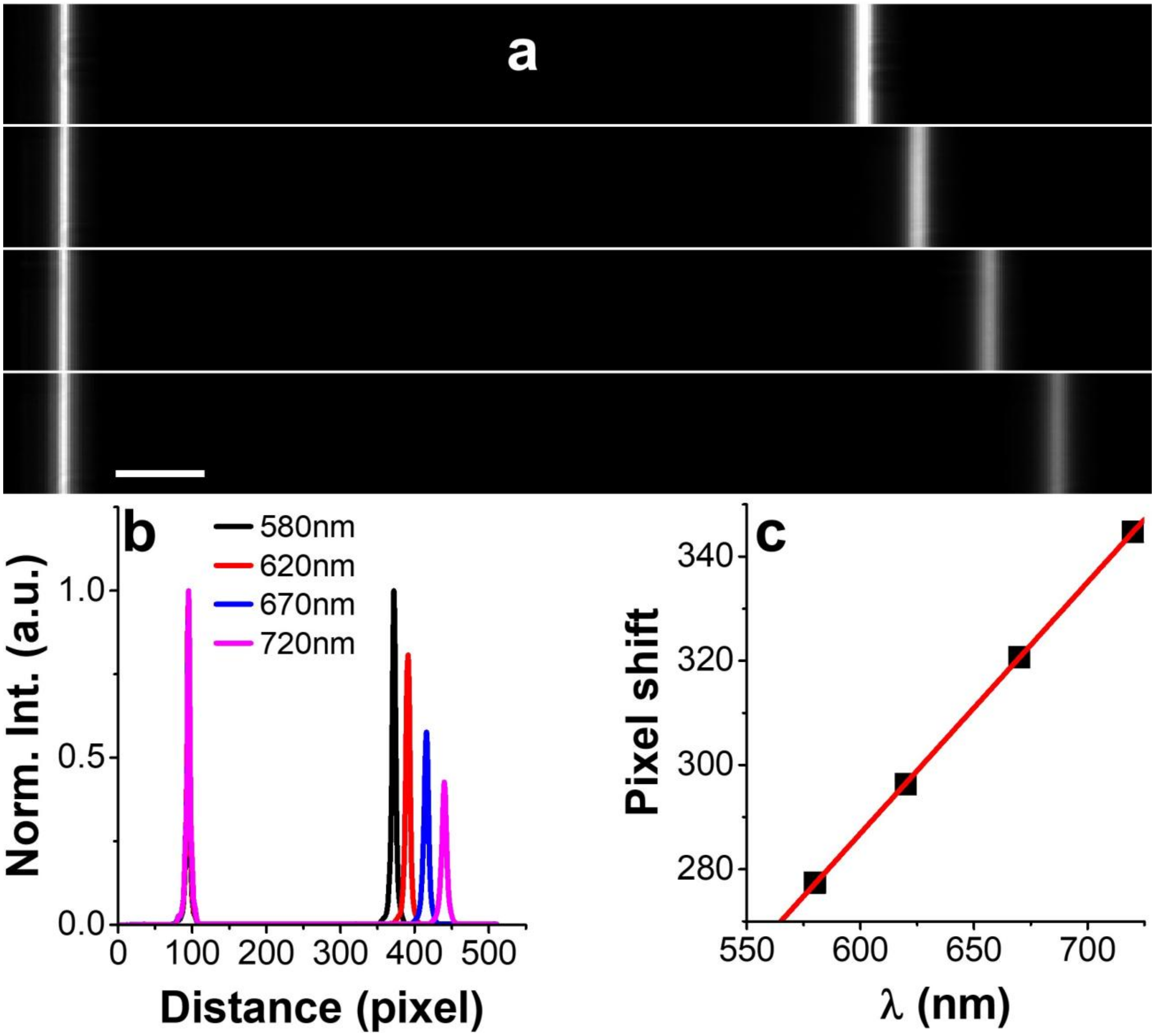

**Figure S5. System calibration for optical grating-based SR-SMLM using narrow band filters.** (a) zero- and first-order image of the optical slit at different wavelengths. (b) Cross-sectional intensity profiles under different wavelengths. (c) Calibration curve of the number of lateral pixel shift over wavelength in first-order image relative to the zero-order image. Scale bar: 5 μm.

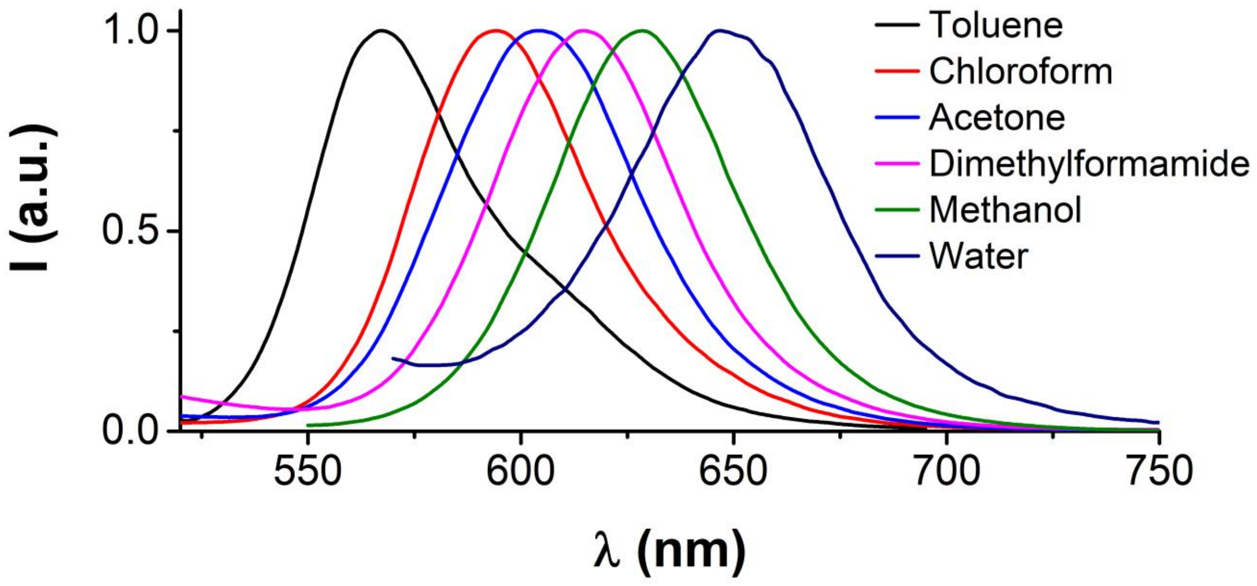

**Figure S6. Dependence of emission spectrum of Nile red on solvent polarity.**

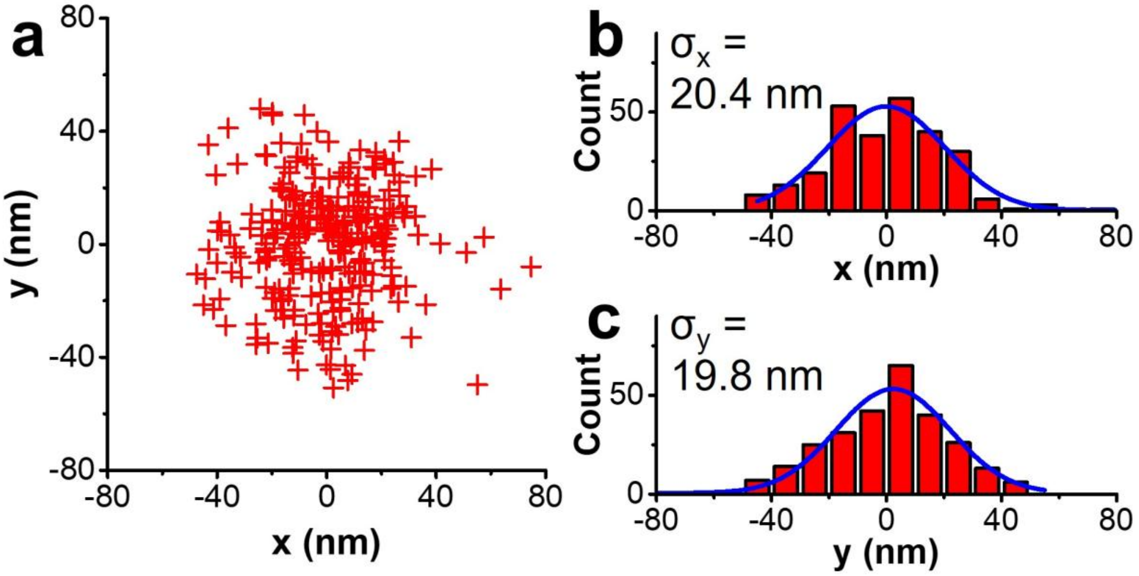

**Figure S7. Localization precision of SP-STORM for Nile red dye.** (a) Cluster analysis of locations. (b, c) Fitting histogram distributions in x, y gives standard deviation of $\sigma_x$ = 20.4 and $\sigma_y$ = 19.8 nm respectively, given 2D spatial resolution ~47 nm (FWHM).

**S9. Captions to Supporting Movies**

**Supporting Movie 1:** The simultaneously recorded single-molecule images in transformed and unmodified channels for SP-STORM imaging of Nile red in live cells, which was obtained within 5 s for the sample shown in Fig. 3a. The EMCCD camera recorded at 200 frames per second (FPS), but the movie is played at the video frame rate of 50 FPS. Scale bar: 5 µm.

**Supporting Movie 2:** SP-STORM imaging of Nile red stained endoplasmic reticulum with 15-sec temporal resolution. The movie is played at 4 frames per second (FPS). Scale bar: 2 µm.
